# Local keratinocyte-nociceptor interactions enhance obesity-mediated small fiber neuropathy via NGF-TrkA-PI3K signaling axis

**DOI:** 10.1101/2024.07.12.603316

**Authors:** Yuta Koui, Shuxuan Song, Xinzhong Dong, Yoh-suke Mukouyama

## Abstract

The pathology of diabetic small fiber neuropathy, characterized by neuropathic pain and axon degeneration, develops locally within the skin during the stages of obesity and pre-diabetes. However, the initiation and progression of morphological and functional abnormalities in skin sensory nerves remains elusive. To address this, we utilized ear skin from mice with diet-induced obesity (DIO), the mouse models for obesity and pre-type 2 diabetes. We evaluated pain-associated wiping behavior and conducted *ex vivo* live Ca^2+^ imaging of the DIO ear skin to detect sensory hypersensitivity. Our findings reveal sensory hypersensitivity in skin nociceptive axons followed by axon degeneration. Further mechanistic analysis identified keratinocytes as a major source of nerve growth factor (NGF) in DIO skin, which locally sensitizes nociceptors through NGF-mediated signaling. Indeed, the local inactivation of NGF and its receptor TrkA-mediated downstream signaling, including the phosphoinositide 3-kinases (PI3K) pathway, suppresses sensory hypersensitivity in DIO skin. Thus, targeting these local interactions between keratinocytes and nociceptors offers a therapeutic strategy for managing neuropathic pain, avoiding the adverse effects associated with systemic interventions.

## Introduction

Patients with diabetes often develop neuropathic pain in the skin of peripheral tissues such as legs and hands in the early stage of diabetes including obesity or prediabetes, called small fiber neuropathy ^1–3^. The primary symptoms of small fiber neuropathy include burning or lancinating pain in the skin ^4,5^ and a reduction in intraepidermal nerve fibers ^6^, often accompanied by abnormal dermal capillaries ^7–9^. Considering that the etiology and pathogenesis of small fiber neuropathy are presently thought to be influenced by systemic hyperglycemia, dyslipidemia, and insulin resistance, the precise mechanism through which local aberrant signals impact the pathophysiological processes of small fiber neuropathy in the skin remains unclear.

Previous studies employing diabetic mouse models successfully recapitulated the pathologies of small fiber neuropathy, including mechanical allodynia and axon degeneration in the footpad skin, abnormal hypersensitivity in sensory neurons of dorsal root ganglions (DRGs), and vascular abnormalities in sciatic nerves ^7,9–11^. However, the mechanisms underlying the initiation and progression of morphological and functional abnormalities in sensory axons and capillaries within the skin of the diabetic mouse models are still inconclusive. Since neuropathic pain and small fiber degeneration are thought to occur during the stages of obesity and pre-diabetes ^12–14^, we employed mouse ear skin to address the underlying mechanisms of small fiber neuropathy in mice with diet-induced obesity (DIO), the mouse models for obesity and pre-type 2 diabetes. We developed a new behavioral assay in which capsaicin application elicits pain-associated wiping behavior of the ears. This was combined with primary sensory neuron-specific GCaMP3 Ca^2+^ imaging to detect hypersensitivity of peripheral terminals of nociceptive neurons in the skin of the ears. Our studies not only reveal the pathogenesis of small fiber neuropathy in the DIO ear skin but also uncover the dominant role of keratinocyte-derived nerve growth factor (NGF) within the epidermis in sensitizing peripheral terminals of nociceptive neurons through NGF-TrkA-PI3K signaling axis.

## Results

### Diet-induced obesity enhances neuropathic pain behavior in mice

To initially assess the link between neuropathic pain behavior and hypersensitivity of peripheral terminals of nociceptive neurons in the skin of mice with obesity or pre-diabetes, we chose to utilize the ear skin of DIO mice due to its simple structure, thin layer, and high accessibility. Mice were fed a high-fat diet from 6 to 30 weeks-of-age to develop DIO (Fig. 1a). Capsaicin, a ligand for transient receptor potential vanilloid subtype 1 (TRPV1) channel, was applied to the skin behind the ears (the outer layer of the ear skin) of both control and DIO mice (Fig. 1b). Since the noxious pain stimuli evoke forelimb wiping responses ^15,16^, we counted those responses for 600 seconds after the capsaicin application (Fig 1b). Mice with a high-fat diet exhibited a significant increase in body weight from 18 weeks-of-age onward (Fig. 1c). Blood glucose and serum insulin levels gradually increased from 18 weeks-of-age and were significantly upregulated from 22 weeks-of-age onward (Fig. 1d, e). Despite the capsaicin application evoking forelimb wiping responses at the stimulus site of ear skin in both control and DIO mice, the wiping bouts were significantly increased in DIO mice at 22 weeks-of-age (Fig. 1f-l). At 30 weeks-of-age, the wiping bouts were slightly enhanced in DIO mice; however, there was a large dispersion between replicates and no significant difference was observed between control and DIO mice (Fig. 1j-l). Thus, the capsaicin-mediated neuropathic pain behaviors in DIO mice peaked at 22 weeks-of-age, after which these pain behaviors were subsequently attenuated by the time they reached 30 weeks-of-age.

**Fig. 1.**
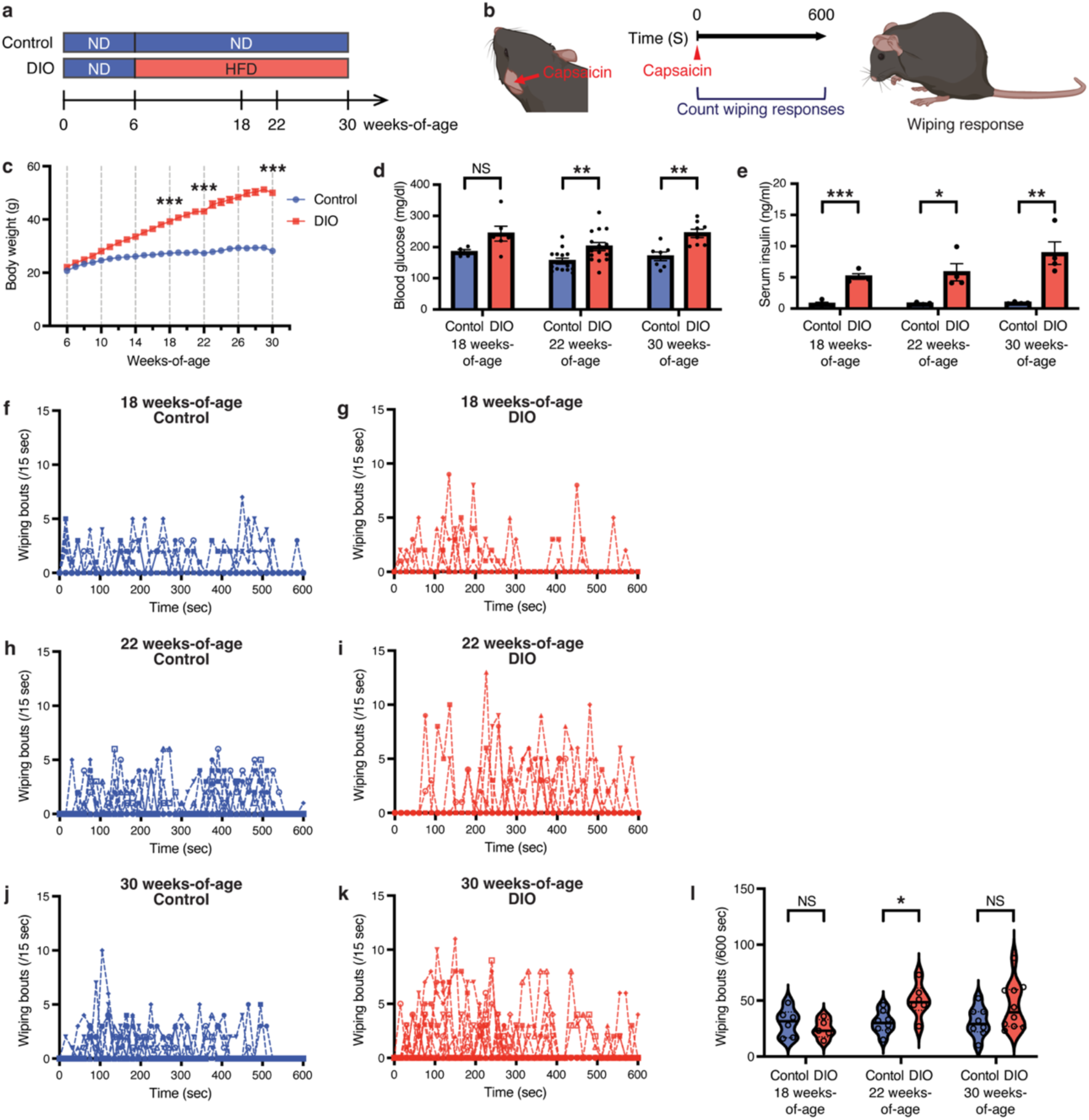
DIO enhances neuropathic pain behavior in mice. (**a**) Schematic diagram of the induction of diet-induced obesity (DIO) in mice. Mice were fed either a high-fat diet (HFD) to induce DIO or a normal diet (ND) for control from 6 to 30 weeks-of-age. (**b**) Illustrations depicting the capsaicin-mediated acute pain behavior assay in mouse ear skin: Capsaicin was applied to the skin behind the ears (the outer layer of the ear skin) of both control and DIO mice. Forelimb wiping responses were recorded for 600 seconds following capsaicin application. (**c**) Body weights of control (N=33) and DIO (N=38) mice from 6 to 30 weeks-of-age. (**d**) Blood glucose levels for both control and DIO mice at 18, 22, and 30 weeks-of-age (N=4, 14, 7 control; N=6, 15, 8 DIO at 18, 22, and 30 weeks-of-age, respectively). (**e**) Serum insulin levels for both control and DIO mice at 18, 22, and 30 weeks-of-age (N=4, 4, 4 control; N=4, 4, 4 DIO at 18, 22, and 30 weeks-of-age, respectively). (**f**-**k**) Forelimb wiping responses following capsaicin application in both control (f, h, j) and DIO (g, i, k) mice at 18 weeks-of-age (f, N=6 control; g, N=5 DIO), 22 weeks-of-age (h, N=8 control; i, N=6 DIO), and 30 weeks-of-age (j, N=8 control; k, N=10 DIO). Each dot represents wiping bouts per 15 seconds. (**l**) Violin plots showing total forelimb wiping responses following capsaicin application in both control and DIO mice at 18, 22, and 30 weeks-of-age. Results are shown as the mean ± SEM (C, D, and E). *p<0.05, **p<0.01, ***p<0.001; NS, not significant (p > 0.05). *P* values were determined by the parametric two-tailed *t* test. The schematic diagrams were partially created with BioRender.com.

### Diet-induced obesity triggers sensory hypersensitivity before skin axon degeneration occurs

We next measured the sensitivity of peripheral terminals of nociceptive neurons in the epidermis of the ear skin using *Pirt-GCaMP3* calcium indicator mice, in which *GCaMP3* is driven by the sensory neuron-specific *Pirt* promoter ^17^. We dissected the outer layer of the ear skin from *Pirt-GCaMP3* mice for *ex vivo* live Pirt-GCaMP3 Ca^2+^ imaging of the ear skin explants (Supplementary Fig. 1a). We conducted imaging of branched sensory axon and terminal activation in response to 2 µM and 10 µM capsaicin applications, based on GCaMP3 fluorescence within the center region of the ear skin epidermis at a depth of 20 μm (Supplementary Fig. 1a, b). After capturing time-lapse Pirt-GCaMP3 Ca^2+^ images for 600 seconds, individual axons were selected from each image for quantification: Ca^2+^ transient (ΔF/F_0_) in each selected axon and the area under the curve (AUC) for the integrated Ca^2+^ elevation were then quantified (Supplementary Fig. 1c-e). At 18 weeks-of-age, 2 μM capsaicin application did not show any significant difference in Ca^2+^ responses in sensory axons within the ear skin explants between control and DIO mice. However, 10 μM capsaicin application evoked Ca^2+^ responses in the ear skin explants from DIO mice compared to controls (Fig. 2a-d and Supplementary Fig. 2a-d). These results suggest that sensory hypersensitivity is not fully developed in DIO skin at 18 weeks-of-age. At 22 weeks-of-age, both 2 µM and 10 μM capsaicin applications evoked Ca^2+^ responses in the ear skin explants from DIO mice compared to controls, suggesting that sensory hypersensitivity is fully developed in DIO ear skin at 22 weeks-of-age (Fig. 2f-i and Supplementary Fig. 2e-h). At 30 weeks-of-age, 2 μM capsaicin application did not show any significant difference in Ca^2+^ responses in the ear skin explants between control and DIO mice. Conversely, 10 μM capsaicin applications attenuated Ca^2+^ responses in the ear skin explants from DIO mice compared to controls (Fig. 2k-n and Supplementary Fig. 2i-l). Collectively, the *ex vivo* live Pirt-GCaMP3 Ca^2+^ imaging of the ear skin explants from DIO mice demonstrated that the capsaicin-mediated sensory hypersensitivity peaked at 22 weeks-of-age, followed by a subsequent attenuation at 30 weeks-of-age. This hypersensitivity in the epidermis of the ear skin corresponds to the forementioned capsaicin-mediated neuropathic pain behaviors.

**Fig. 2.**
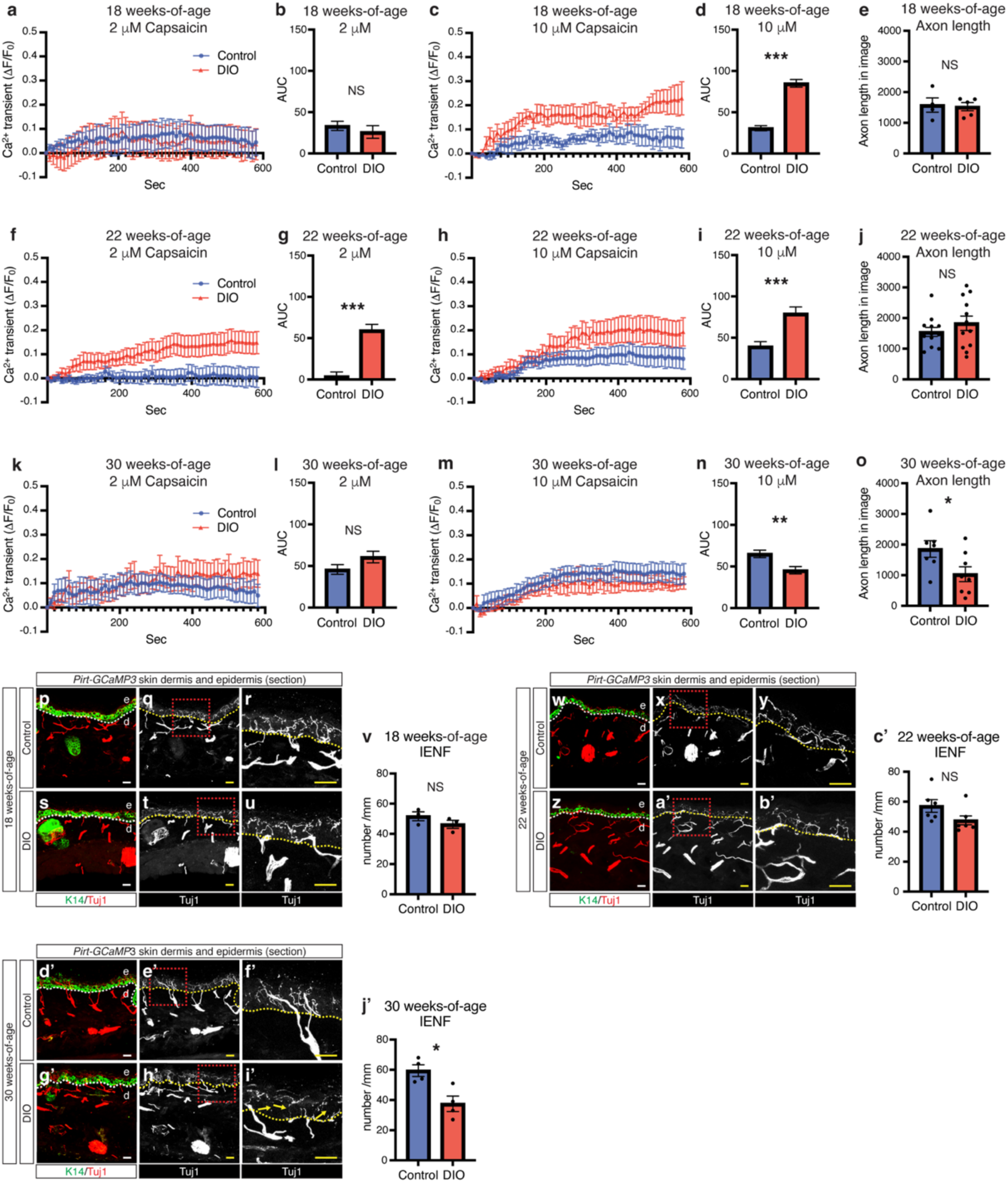
DIO induces sensory hypersensitivity preceding skin axon degeneration within the epidermis. (**a-d**, **f-i**, and **k-n**) Quantification of Ca^2+^ responses within the ear skin explants from *Pirt-GCaMP3* control (blue) and DIO (red) mice at 18 weeks-of-age (a-d, N=4 control, N=5 DIO), 22 weeks-of-age (f-i, N=12 control, N=12 DIO), and 30 weeks-of-age (k-n, N=6 control, N=7 DIO). The amplitude of the Ca^2+^ transient was evoked by 2 μM capsaicin (a, f, and k) and 10 μM capsaicin (c, h, and m). Ca transient was normalized by baseline Ca^2+^ transient (ΔF/F_0_). The integrated Ca^2+^ transient (ΔF/F_0_) was calculated as the area under curve (AUC) (b, d, g, i, l, and n). (**e**, **j** and **o**) Total length of GCaMP3-expressing axons in the epidermis at 18 weeks-of-age (e, N=4 control, N=5 DIO), 22 weeks-of-age (j, N=12 control, N=12 DIO), and 30 weeks-of-age (o, N=7 control, N=8 DIO). (**p**-**u**, **w**-**b’**, and **d’-i’**) Representative images of double-immunofluorescence staining of ear skin sections of *Pirt-GCaMP3* control (p-r, w-y, and d’-f’) or DIO (s-u, z-b’, and g’-i’) mice at 18, 22, and 30 weeks-of-age. Antibodies to the keratinocyte marker K14 (green) and the pan-axon/neuron marker neuron-specific class III ß-tubulin (Tuj1, red or white) were used. The dotted box regions in (q), (t), (x), (a’), (e’), and (h’) were magnified in (r), (u), (y), (b’), (f’), and (i’), respectively. Dashed lines indicate the border between the epidermis (e) and the dermis (d). Yellow arrows in (i’) indicate degenerated axons in the epidermis. Scale bars: 20 μm. (**v**, **c’**, and **j’**) Quantification of intraepidermal nerve fibers (IENF) in each image at 18 weeks-of-age (v, N=3 control, N=3 DIO), 22 weeks-of-age (c’, N=6 control, N=6 DIO), and 30 weeks-of-age (j’, N=4 control, N=4 DIO). The linear IENF density is calculated and expressed as the number of fibers per millimeter of epidermal length (IENF/mm). Results are shown as the mean ± SEM. *p<0.05, **p<0.01, ***p<0.001; NS, not significant (p > 0.05). *P* values were determined by the parametric two-tailed *t* test.

The observation that hypersensitivity was attenuated at 30 weeks-of-age, along with the reduction in neuropathic pain behaviors, prompted us to examine whether these attenuations result from axon degeneration in nociceptive neurons within the epidermis of the ear skin. Indeed, Pirt-GCaMP3 Ca^2+^ imaging at 30 weeks-of-age indicated a reduction in axon length within the epidermis of DIO ear skin compared to controls (Fig. 2o), whereas at both 18 and 22 weeks-of-age, there was no significant difference observed in axon length between control and DIO ear skin (Fig. 2e, j). Similarly, our section immunohistochemical analysis of control and DIO ear skin clearly showed a significant reduction in intraepidermal nerve fiber (IENF) density, quantified by the number of nerve fibers crossing the epidermal-dermal junction, in DIO ear skin at 30 weeks-of-age (Fig. 2d’-j’), whereas at both 18 and 22 weeks-of-age, there was no significant difference in the IENF density between control and DIO ear skin (Fig. 2p-c’). Collectively, these data suggest that DIO ear skin develops sensory hypersensitivity before axon degeneration.

We further examined morphological changes in sensory nerves and blood vessels in DIO ear skin (Supplementary Fig. 3a, b). Previously, we demonstrated the co-branching of sensory nerves and large-diameter blood vessels covered by vascular smooth muscle cells (VSMCs) in the deep dermis of ear skin ^18^. At 22 and 30 weeks-of-age, no significant changes were observed in the branching patterns of sensory nerves and blood vessels in the deep dermis between control and DIO ear skin (Supplementary Fig. 4a-t). Additionally, there were no significant changes found in VSMC coverage of large-diameter blood vessels (Supplementary Fig. 4a-t) or myelination of sensory nerves (Supplementary Fig. 4p’-u’) between control and DIO ear skin. In contrast, given that skin capillary endothelial cells form non-fenestrated blood vessels with adhesion junctions between endothelial cells ^19^, we observed increased expression of the vascular hyperpermeability marker, plasmalemma vesicle-associated protein (PLVAP) ^20^, in DIO skin vasculature adjacent to the boundary between the epidermis and dermis at 22 and 30 weeks-of-age, but not 18 weeks-of-age (Supplementary Fig. 4u-o’). These data suggest a correlation between the emergence of increased vascular permeability in the skin vasculature and the development of sensory hypersensitivity in DIO mice. Given that immune cells are known to modulate nociceptor terminals in the pathological skin ^21^, we observed no significant change in the number of CD45^+^ immune cells within the epidermis of DIO ear skin, where hypersensitivity in sensory axon terminal was detected (Supplementary Fig. 4v’-b’’).

### Diet-induced obesity enhances the expression of NGF, sensitization of nociceptors, in the epidermis

Neurotrophic factors support the growth, differentiation, and maintenance of neuronal cells in development, but they also contribute to neurological disorders in both peripheral and central nervous systems ^22^. In chronic diseases, among those neurotrophic factors, NGF is a known to promote pain reactions in sensory nerves by inducing TRPV1 expression, facilitating TRPV1 trafficking to cell membrane, or sensitizing TRPV1 receptor ^23–27^. Moreover, single injections of NGF into the healthy human skin evoked pain ^28,29^. To address whether NGF affects sensitization of peripheral terminals of nociceptive neurons in the epidermis of ear skin, we initially examined NGF expression in the epidermis from control and DIO ear skin at 22 weeks-of-age. Among the three neurotrophic factors tested, we observed increased expression of *Ngf* in the epidermis of DIO ear skin compared to control, whereas no significant changes were detected in the expression levels of *Bdnf* and *Ntf3* (Fig. 3a). In support of this result, Pirt-GCaMP3^+^ sensory nerves in the epidermis express NGF receptor, TrkA, but not TrkB and TrkC (Supplementary Fig. 5a-i). We next addressed whether the onset of NGF expression in the epidermis correlates with the development of sensory hypersensitivity in DIO mice (Fig. 3b). At 22 weeks-of-age, X-gal staining of ear skin sections from *NGF-LacZ* mice showed increased expression of NGF in the epidermis of DIO ear skin compared to control (Fig. 3c, d). For quantification measurements, we further validated the enhanced NGF expression in the epidermis of DIO ear skin using a fluorescent version of LacZ detection, SPiDER-βgal staining in conjunction with the keratinocyte markers K10 and K14 (Fig. 3e-r). In contrast, at 18 weeks-of-age, we did not observe any significant difference in NGF expression in the epidermis between control and DIO ear skin (Supplementary Fig. 6a-o, Fig. 3s). At 30 weeks-of-age, there was a significant difference in NGF expression in the epidermis between control and DIO ear skin, albeit with a reduced expression level compared to that at 22 weeks-of-age (Supplementary Fig. 7a-o, Fig. 3s). We should note that dermal NGF expression was primary detected in VSMCs of large-diameter blood vessels, as observed through whole-mount X-gal or SPiDER-βgal staining. However, no significant differences were observed in the expression levels between control and DIO ear skin, nor between those different stages (Supplementary Fig. 5j-q, 6p-w, 7p-w). Collectively, these data suggest a correlation between the onset of NGF expression in the epidermis of DIO ear skin and the development of sensory hypersensitivity.

**Fig. 3.**
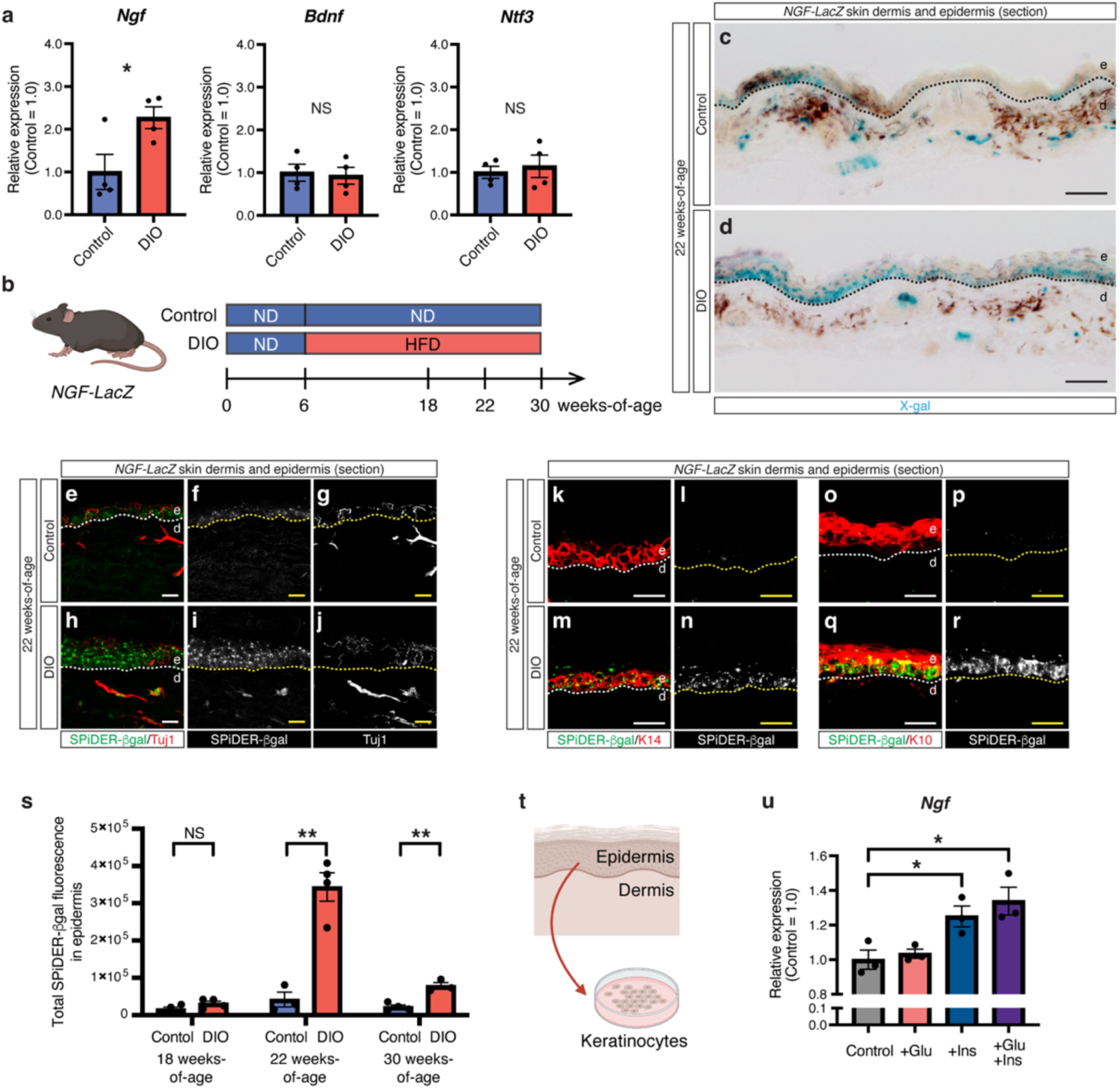
DIO enhances NGF expression in the epidermis. (**a**) Relative mRNA expression levels of neurotrophic factors (*Ngf*, *Bdnf*, and *Ntf3*) in the epidermis of ear skin from control and DIO mice at 22 weeks-of-age. Expression levels are normalized by those in the epidermis of ear skin from control mice. N=4 in each group. (**b**) Schematic diagram of the induction of DIO in *NGF-LacZ* mice. (**c**-**d**) Representative images of X-gal staining of ear skin sections of *NGF-LacZ* control (c) and DIO mouse (d) at 22 weeks-of-age (blue). Scale bars: 100 μm. (**e**-**r**) Representative double immunofluorescence images of ear skin sections of *NGF-LacZ* control (e-g, k-l, and o-p) and DIO mice (h-j, m-n, and q-r) at 22 weeks-of-age. SPiDER-βgal (e, h, k, m, o, and q, green; f, i, l, n, p, and r, white) together with antibodies to Tuj1 (e and h, red; g and j, white), K14 (k and m, red), or K10 (o and q, red) were used. Dashed lines (c-d, e-r) indicate the border between the epidermis (e) and the dermis (d). (**s**) Total SPiDER-βgal fluorescence in the epidermis of ear skin from *NGF-LacZ* mice at 18 weeks-of-age (N=4 control, N=4 DIO), 22 weeks-of-age (N=3 control, N=4 DIO), and 30 weeks-of-age (N=4 control, N=3 DIO). See also fig. S6 and S7. (**t**) Schematic diagram of primary mouse keratinocyte culture. Keratinocytes were isolated from adult ear skin and cultured with glucose, insulin, or a combination of both (**u**) Relative *Ngf* mRNA expression levels in primary mouse keratinocytes cultured with glucose (+Glu), insulin (+Ins), or a combination of both (+Glu +Ins). Expression levels are normalized by those in primary mouse keratinocytes cultured without glucose and insulin. N=3 in each group. Results are shown as the mean ± SEM. *p<0.05, **p<0.01; NS, not significant (p > 0.05). *P* values were determined by the parametric two-tailed *t* test. The schematic diagrams were partially created with BioRender.com.

We then examined what controls NGF expression in the epidermis. Considering that keratinocytes constitute over 95% of the cells in the epidermis, we initially isolated keratinocytes from ear skin of adult mice, cultured them with glucose, insulin, or a combination of both, and subsequently assessed *Ngf* expression (Fig. 3t, u). A high level of insulin enhances *Ngf* expression in primary keratinocytes in culture, whereas a high level of glucose did not show any inductive effect on its expression (Fig. 3u). These data suggest that epidermal keratinocytes are a major source of NGF in response to increased levels of insulin in DIO ear skin.

### Inhibition of NGF-TrkA-PI3K signaling suppresses sensory hypersensitivity in the skin of diet-induced obesity mice

To address whether NGF-TrkA signaling is necessary for sensitization of peripheral terminals of nociceptive neurons in the epidermis of DIO ear skin, we pre-treated the ear skin explants from *Pirt-GCaMP3* DIO mice at 22 weeks-of-age with an anti-NGF neutralizing antibody (Fig. 4a, b). Subsequently, we conducted Pirt-GCaMP3 Ca^2+^ imaging to assess sensory axon and terminal activation in response capsaicin applications (Fig. 4b). The capsaicin-mediated hypersensitivity observed in DIO ear skin explants was significantly suppressed by pre-treating the explants with the anti-NGF neutralizing antibody (Fig. 4c-f, Supplementary Fig. 8a-d). These data clearly suggest that NGF-TrkA signaling is necessary for capsaicin-mediated hypersensitivity in the epidermis of DIO ear skin (Supplementary Fig. 9). Among the downstream targets of NGF-TrkA signaling, such as phosphoinositide 3-kinases (PI3K), mitogen-activated protein kinase (MAPK), and phospholipase C (PLC), which are known to be involved in nociceptor sensitization through the capsaicin receptor TRPV1 ^27,30,31^, we initially examined whether inhibiting PI3K leads to the blockage of capsaicin-mediated hypersensitivity. To address this, we pre-treated the ear skin from *Pirt-GCaMP3* DIO mice at 22 weeks-of-age with Wortmannin, a PI3K inhibitor (Fig. 4a, b). Like the anti-NGF neutralizing antibody treatment, the capsaicin-mediated hypersensitivity observed in DIO ear skin explants was significantly suppressed by pre-treating the explants with Wortmannin (Fig. 4g-j, Supplementary Fig. 8e-h). These data suggest that NGF-TrkA-PI3K signaling is necessary for capsaicin-mediated hypersensitivity in the epidermis of DIO ear skin (Supplementary Fig. 9). Moreover, this procedure is able to screen small compound inhibitors to suppress the capsaicin-mediated hypersensitivity observed in DIO ear skin explants.

**Fig. 4.**
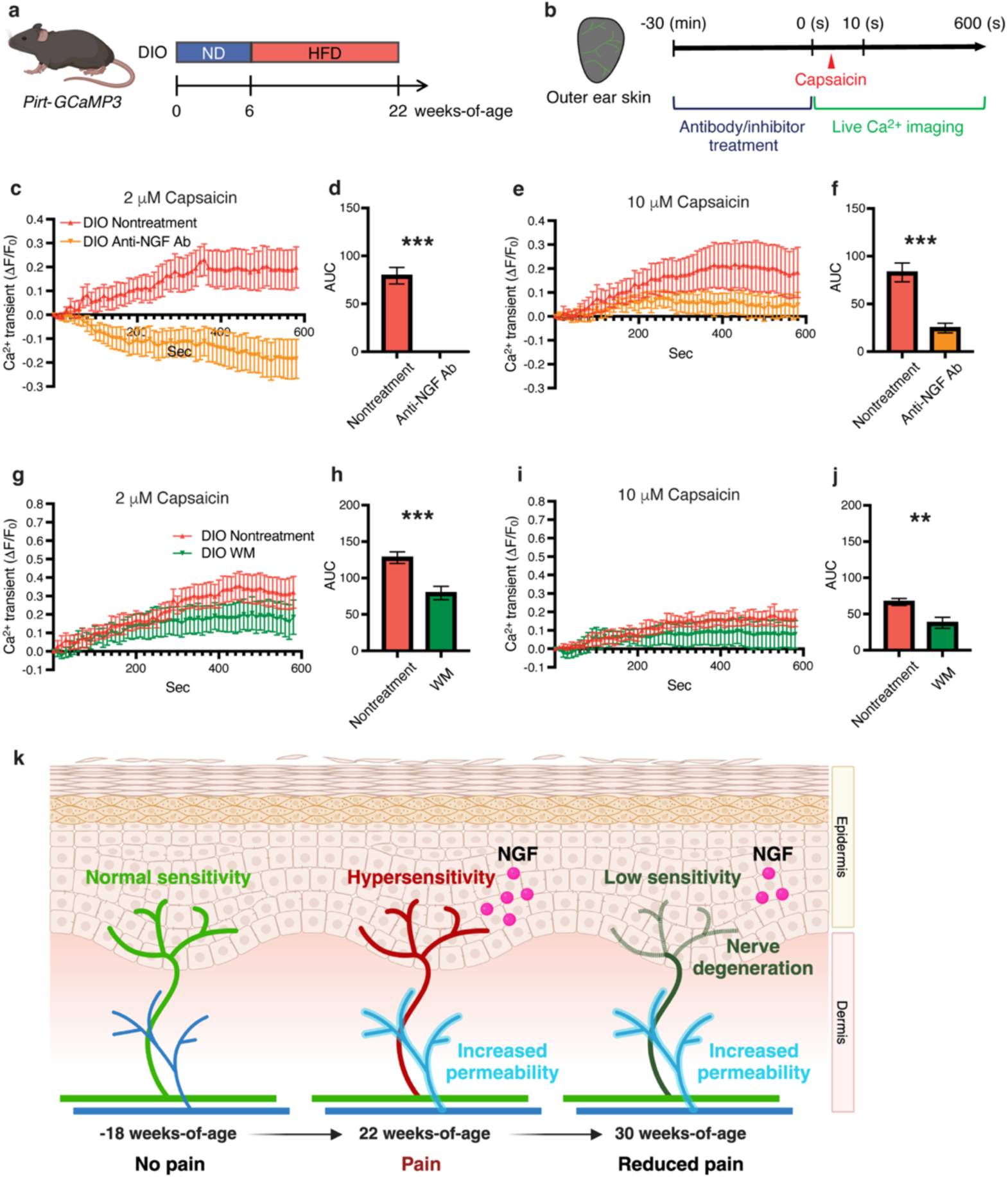
Inhibition of NGF-TrkA-PI3K signaling suppress hypersensitivity in DIO skin. (**a**) Schematic diagram of the induction of DIO in *Pirt-GCaMP3* mice. (**b**) Schematic diagram illustrating *ex vivo* live Pirt-GCaMP3 Ca^2+^ imaging of ear skin explants treated with anti-NGF neutralizing antibody or Wortmannin, a PI3K inhibitor. The ear skin explants from *Pirt-GCaMP3* DIO mice at 22 weeks-of-age were pre-treated with either anti-NGF neutralizing antibody or Wortmannin for 30 min prior to Pirt-GCaMP3 Ca^2+^ imaging. (**c-j**) Quantification of Ca^2+^ responses within the ear skin explants treated with or without anti-NGF neutralized antibody treatment (c-f, Anti-NGF Ab) or Wortmannin treatment (g-j, WM). N=6 in each group. The amplitude of the Ca^2+^ transient was evoked by 2 μM capsaicin (c and g) and 10 μM capsaicin (e and i). Ca transient is normalized by baseline Ca transient (ΔF/F_0_). The integrated Ca^2+^ transient (ΔF/F_0_) was calculated as AUC (d, f, h, and j). Results are shown as the mean ± SEM. **p<0.01, ***p<0.001. *P* values were determined by the parametric two-tailed *t* test. (**k**) Graphical summary for neuropathic pain behaviors, sensory hypersensitivity, epidermal axon projections, vascular permeability, and epidermal NGF expression in the ear skin of DIO mice at 18, 22, and 30 weeks-of-age. The schematic diagrams and graphic summary were partially created with BioRender.com.

## Discussion

We present evidence supporting the role of keratinocyte-derived NGF as a local signal sensitizing the peripheral terminals of nociceptive neurons through NGF-TrkA-PI3K signaling within the epidermis of DIO ear skin, thereby augmenting neuropathic pain behaviors (Fig. 4k). At mechanistic level, elevated insulin levels enhance *Ngf* expression in primary keratinocytes from the epidermis. Although no significant changes were observed in serum insulin levels in DIO mice between 18 and 22 weeks-of-age, insulin levels appear to rise sufficiently by 22 weeks-of-age to induce *Ngf* expression in keratinocytes within the non-capillary epidermis. Considering the increased vascular permeability in DIO ear skin vasculature at 22 weeks-of-age, but not at 18 weeks-of-age, one scenario is that the increased vascular permeability may facilitate the diffusion of circulating insulin from dermal capillaries into the non-capillary epidermis in DIO ear skin. In this scenario, reducing or blocking vascular permeability could regulate insulin levels in the epidermis of DIO ear skin, potentially leading to decreased epidermal NGF expression and consequently attenuating sensory hypersensitivity.

The observation that DIO ear skin undergoes axon degeneration in nociceptive neurons within the epidermis at 30 weeks-of-age raises the intriguing question of what triggers this degeneration. Previous studies have demonstrated that prolonged activation of nociceptive neurons accelerates axon degeneration in DIO skin ^10^. However, the precise mechanisms driving axon degeneration due to prolonged activation of nociceptive neurons remain incompletely elucidated. In this context, inhibiting the sensitization of nociceptors through targeting local NGF-TrkA-PI3K signaling could potentially prevent axon degeneration in DIO skin. It is important to note that the progressive degeneration of epidermal nerve fibers of nociceptive neurons may lead to damage of terminal arbors of myelinated peripheral sensory nerves in the deep dermis, indicating a pathology of large fiber neuropathy generally occurring at a late stage of diabetes. Indeed, some patients with small fiber neuropathy develop large fiber neuropathy with distal numbness over time as their diabetes progress ^32^.

Considering the close proximity of keratinocytes to sensory terminals within the epidermis, it is established that keratinocytes contribute to mechanical, cold and heat sensation by detecting these stimuli and releasing neuroactive molecules for sensory neurons ^33–35^. In line with this concept, we present evidence that the crosstalk between keratinocytes and nociceptor terminals plays a crucial role in neuropathic pain through NGF-TrkA-PI3K signaling. Our understanding of the local interactions between keratinocytes and nociceptor terminals within the epidermis, along with the identification of potential target signaling pathways such as NGF-TrkA-PI3K signaling to regulate nociceptor sensitization, could serve as the foundation for novel therapeutic strategies to manage neuropathic pain in diabetic small fiber neuropathy. Importantly, these approaches could be topically applied, potentially circumventing drug side effects associated with systemic interventions.

## Methods

### Mice

All animal procedures were approved by the National Heart, Lung, and Blood institute (NHLBI) Animal Care and Use Committee in accordance with National Institutes of Health (NIH) research guidelines for the care and use laboratory animals. The following mice were used in this study: C57BL/6J mice (The Jackson Laboratory), *Pirt-GCaMP3* mice ^17^, and *NGF-LacZ* mice ^36^. C57BL/6J, *Pirt-GCaMP3* heterozygous, and *NGF-LacZ* heterozygous male mice were randomly separated into two groups and fed either a high-fat diet (60% fat, 20% protein, and 20% carbohydrate kcal; Research Diets, D12492) to induce DIO or a normal diet (10% fat, 20% protein, and 70% carbohydrate kcal) for control from 6 weeks-of-age with free access to water.

### Body weight, blood glucose, and serum insulin measurement

Body weights of all mice were measured weekly at the same time of the day. Blood glucose levels were measured using a glucose meter, AlphaTrak 3 blood glucose monitoring system (Zoetis) after 4-6 hours fasting. Blood samples were collected after cardiac puncture. Serum insulin levels were measured by Mouse Ultrasensitive Insulin ELISA system (ALPCO) according to manufacturer’s protocol.

### Pain behavior assay

The control or DIO mice were placed in the plastic clear test chamber and habituated for 10 min. The 10 ml capsaicin solution (0.1 mM in ethanol) was applied into the skin behind the ear after habituation. The mouse behavior was videorecorded for 10 min. The wiping responses with a forelimb toward the stimulus site was manually counted based on the recorded videos. Wiping bouts/600 seconds were calculated for comparison between control and DIO mice at each stage.

### Ex vivo live Ca^2+^ imaging of mouse ear skin

*Pirt-GCaMP3* mice at 18, 22, and 30 weeks-of-age were sacrificed by CO_2_ asphyxiation. Ear skin was acutely dissected from *Pirt-GCaMP3* mice, and then the outer ear skin was isolated for Ca^2+^ imaging. Epidermal branched axons located within 20 mm from the surface of hairy skin were imaged. Skin explants were placed in test chamber filled with Synthetic interstitial fluid (SIF) ^37^ and maintained at room temperature. The skin explants were treated with anti-NGF neutralized antibody (0.02 mg/ml, exalpha, L146M) or Wortmannin (1 mM, Sigma, W1628) for 30 min at room temperature before Ca^2+^ imaging. Ca^2+^ imaging was carried out on a Leica TCS SP5 confocal (Leica) with perfusion of SIF. The skin was stimulated by 2 mM or 10 mM Capsaicin (Sigma, M2028) and then imaged for 10 min to measure GCaMP3 fluorescent levels. The 5 Z-stack images were acquired every 10 seconds. For quantification, time-course images were imported into Image J (NIH) to create maximum intensity projection (MIP) images. Three areas with GCaMP3 (green) positive sensory axons were cropped from the MIP images, and averaged Ca^2+^ transients (DF/F0) were calculated. For comparison of sensory activities in response to capsaicin between control and DIO mice, amount of Ca^2+^ transient that was calculated as area under curve was determined.

### Section immunostaining of mouse ear skin

Outer ear skin was dissected from adult mice, fixed with 4% paraformaldehyde/ PBS at 4°C overnight, sunk in 30% sucrose/PBS at 4°C, and then embedded in OCT compound. Tissues were cryosectioned at 20 mm thickness and collected on pre-cleaned slides (Fisher Scientific, 15-188-48). The samples were incubated in blocking buffer (0.2% TritonX-100, 10% heat inactivated goat or donkey serum in PBS), with diluted primary antibodies 4°C overnight. Staining was performed using Mouse anti-Tuj1 (Tubulin b3, TUBB3) antibody (BioLegend, 801202, 1:200), Goat anti-TrkA antibody (R&D Systems, AF1056-SP, 1:100), Goat anti-TrkB antibody (R&D Systems, AF1494-SP, 1:100), and Goat anti-TrkC antibody (R&D Systems, AF1404-SP, 1:100) to detect axons, Chick anti-GFP antibody (abcam, ab13970, 1:300) to detect GCaMP3 protein in sensory axons, Rabbit anti-MBP (Myelin Basic Protein) antibody (abcam, ab40390, 1:200) to detect myelin membrane in peripheral nerves, Rabbit anti-K10 (Keratin 10) antibody (BioLegend, 905403, 1:400) and Guinea pig anti-K14 (Keratin 14) antibody (PROGEN, GP-CK14, 1:100) to detect keratinocytes, Armenian hamster anti-PECAM-1 antibody (Millipore Sigma, MAB1398Z, 1:300) and Rat anti-PLVAP (BD Biosciences, 553849, 1:100) to detect endothelial cells, and Rat anti-CD45 antibody (eBioscience, 140451-85, 1:100) to detect hematopoietic cells. For immunofluorescent detection, the samples were incubated in blocking buffer containing either Alexa-488-, Alexa-568-, or Alexa-647-conjugated secondary antibodies (Jackson ImmunoResearch or Thermo Fisher Scientific, 1:250). All confocal microscopy was carried out on a Leica TCS SP5 confocal (Leica).

### Measurement of intraepidermal nerve fiber (IENF) density

IENF was quantified by counting the number of sensory axons crossing the epidermal-dermal junction. The linear IENF density was calculated and expressed as the number of fibers per millimeter of epidermal length (IENF/mm) for the comparison between control and DIO mice.

### Measurement of PLVAP^+^ blood vessels

Area of PLVAP^+^ blood vessels were quantified using ImageJ (NIH). The percentage of PLVAP^+^ blood vessels was based on the area of PLVAP^+^ blood vessels within the area of PECAM-1^+^ blood vessels.

### Measurement of CD45^+^ immune cells in the epidermis

The quantity of CD45^+^ immune cells present in the epidermis was counted and expressed as the cell number per millimeter of epidermal length for the comparison between control and DIO mice.

### Whole-mount immunostaining of mouse ear skin

Staining was performed essentially as described previously ^18,38^. After fixation, connective tissues and fat were removed from the dermal side of the ear skin before staining. The samples were incubated in blocking buffer with diluted primary antibodies 4°C overnight. Staining was performed using Alexa-488-conjugated Mouse anti-Tuj1 (Tubulin b3, TUBB3) antibody (BioLegend, 801203, 1:200) to detect axons, Cy3-conjugated Mouse anti-aSMA antibody (Sigma, C6198, 1:250) and Rabbit anti-SM22a antibody (abcam, ab14106, 1:500) to detect smooth muscle cells, and Armenian hamster anti-PECAM-1 antibody (Millipore Sigma, MAB1398Z, 1:300) to detect endothelial cells. For immunofluorescent detection, the samples were incubated in blocking buffer containing either Alexa-488-, Alexa-568-, or Alexa-647-conjugated secondary antibodies (Jackson ImmunoResearch or Thermo Fisher Scientific, 1:250). All confocal microscopy was carried out on a Leica TCS SP5 confocal (Leica).

### Quantitative RT-PCR

Ear skin was incubated in 0.15% Trypsin-EDTA for 45 min at 37°C to separate the epidermis from dermis. Total RNAs from epidermis tissue and primary cultured mouse keratinocytes were extracted using RNeasy Mini kits (QUAGEN) according to the manufacturer’s protocol.

Residual genomic DNA was digested with Ambion DNase I (Thermo Fisher Scientific). First-strand cDNA was synthesized using the SuperScript III First-Strand Synthesis SuperMix (Thermo Fisher Scientific, 18080400). Quantitative RT-PCR was performed using THUNDERBIRD Next SYBR qPCR Mix (TOYOBO). All data of quantitative RT-PCR were calculated using the ddCt method with *Gapdh* as normalization controls. Primers are listed in Table S1.

### X-gal staining of mouse ear skin

Outer ear skin was fixed with 0.25% glutaraldehyde/ PBS for 30 min on ice. After fixation, connective tissues and fat were removed from the dermal side of the ear skin before staining. The tissues were incubated in 1 mg/ml X-gal overnight at 4°C. The tissues were washed with PBS and post-fixed with 4% paraformaldehyde/ PBS. For the section imaging, tissues were further sunk in 30% sucrose/ PBS at 4°C and then embedded in OCT compound. Tissues were cryosectioned at 20 mm thickness.

### SPiDER-ßgal staining of mouse ear skin

Outer ear skin isolated from adult mice was incubated in 20 mM SPiDER-ßgal (Dojindo) for 1 hour at 37°C. Tissues were washed with PBS and then fixed with 4% paraformaldehyde/ PBS for 1 hour on ice. Tissues were further stained together with antibodies as described above.

### Quantification of SPiDER-ßgal signals in epidermis

Total SPiDER-ßgal signals were quantified using ImageJ (NIH). Fluorescent levels were normalized by the area of epidermis in each image.

### Primary mouse keratinocyte culture

Primary mouse keratinocytes were isolated from C57BL/6J male mice according to the previously established protocol ^39^. Isolated mouse keratinocytes were cultured on fibronectin/collagen coated plates at a density of 1,000,000 cells/cm^2^. Cells were replated at day 2 to culture under high-glucose or high-insulin condition for 2 days.

### Statistical analysis

Results were presented as the mean ± SEM. *P* values were determined by the parametric two-tailed *t* test. p < 0.05 was considered statistically significant.

## Data and materials availability

All data needed to evaluate the conclusions are available in the main text or the supplementary materials.

## Acknowledgements

Thanks to T. Clark and the staff of NIH Bldg50 facility for assistance with mouse breeding and care; E. Tyler and A. Hoofring of NIH medical art branch for schematic illustrations; K. Gill for laboratory management, technical support, and manuscript editing; V. Sam and S. Thacker for administrative assistance. Thanks also to members of Laboratory of Stem Cell and Neuro-Vascular Biology for thoughtful discussion and technical help. This work was supported by the Intramural Research Program of the National Heart, Lung, and Blood Institute, National Institutes of Health (HL005702-18 to Y.M.)

## Author contributions

Conceptualization: Y.K., Y.M.; Formal analysis: Y.K., S.S.; Funding acquisition: Y.M.; Investigation: Y.K., S.S.; Methodology: Y.K., S.S.; Project administration: Y.M.; Resources: X.D.; Supervision: Y.M.; Validation: Y.K., S.S.; Visualization: Y.K., S.S.; Writing - original draft: Y.K., Y.M.; Writing - review & editing: Y.K., S.S., Y.M., X.D.

