## Supplementary figures and table for "Local keratinocyte-nociceptor interactions enhance obesity-mediated small fiber neuropathy via NGF-TrkA-PI3K signaling axis"

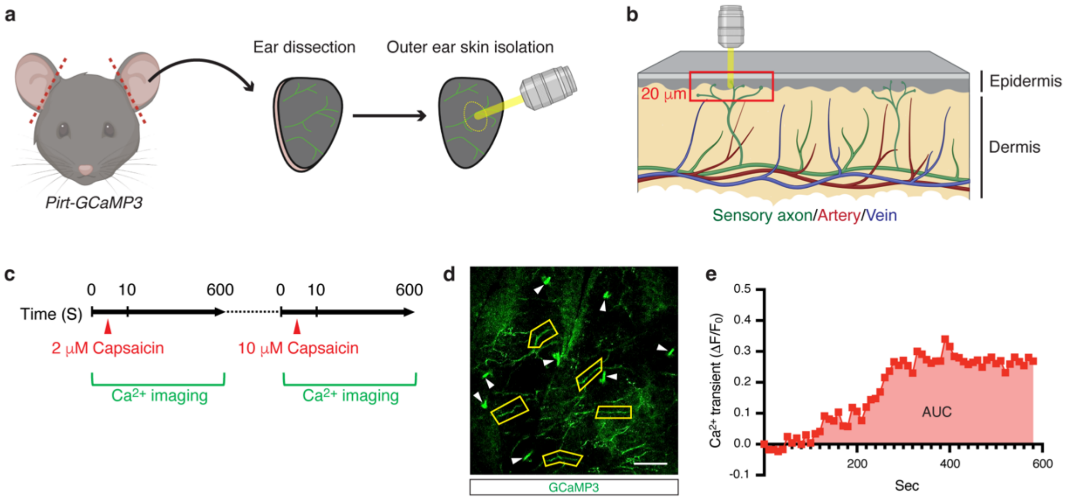


**Supplementary Fig. 1.** ***Ex vivo*** **live Pirt-GCaMP3 $Ca^{2+}$ imaging of ear skin explants.**

(**a**) Illustration depicting the ear skin dissection procedure for *ex vivo* live Pirt-GCaMP3 $Ca^{2+}$ imaging. GCaMP3 fluorescence signals were assessed within the center region of the epidermis of the ear skin (indicated by the dotted circle). (**b**) Illustration of dissected ear skin. GCaMP3 fluorescence signals in branched sensory axons and their terminals within the epidermis of the ear skin at a depth of 20 µm were assessed (indicated by the red boxed region). In the epidermis, sensory axons and their terminals (green) are present, while capillaries are absent; in the dermis, both sensory axons and vasculature, including arteries (red) and veins (blue) are present. (**c**) Schematic diagram illustrating *ex vivo* live Pirt-GCaMP3 $Ca^{2+}$ imaging of ear skin explants. GCaMP3 fluorescence signals were captured for 600 seconds after capsaicin application. There was a brief interval in Pirt-GCaMP3 $Ca^{2+}$ imaging between the ear skin samples treated with 2 µM capsaicin and those treated with 10 µM capsaicin. (**d**) Representative image showing GCaMP3 expression (green) within the epidermis. GCaMP3-expressing axons and their terminals were selected for quantification of fluorescence levels (indicated by the GCaMP3-expressing axons and terminals enclosed by yellow frames). Note that GCaMP3-expressing hair follicles (arrowheads) were excluded from this analysis. Scale bar: 100 µm. (**e**) Representative data showing the time course of the amplitude of the $Ca^{2+}$ transient ($\Delta F/F_0$) evoked by capsaicin application in ear skin explants. $Ca^{2+}$ transient is normalized by baseline $Ca^{2+}$ transient ($\Delta F/F_0$). Each dot represents the average $Ca^{2+}$ transient ($\Delta F/F_0$) from three selected GCaMP3-expressing axons and terminals within the epidermis. The integrated $Ca^{2+}$ transient ($\Delta F/F_0$) was calculated as the area under curve (AUC). The illustrations and schematic diagram were partially created with BioRender.com.

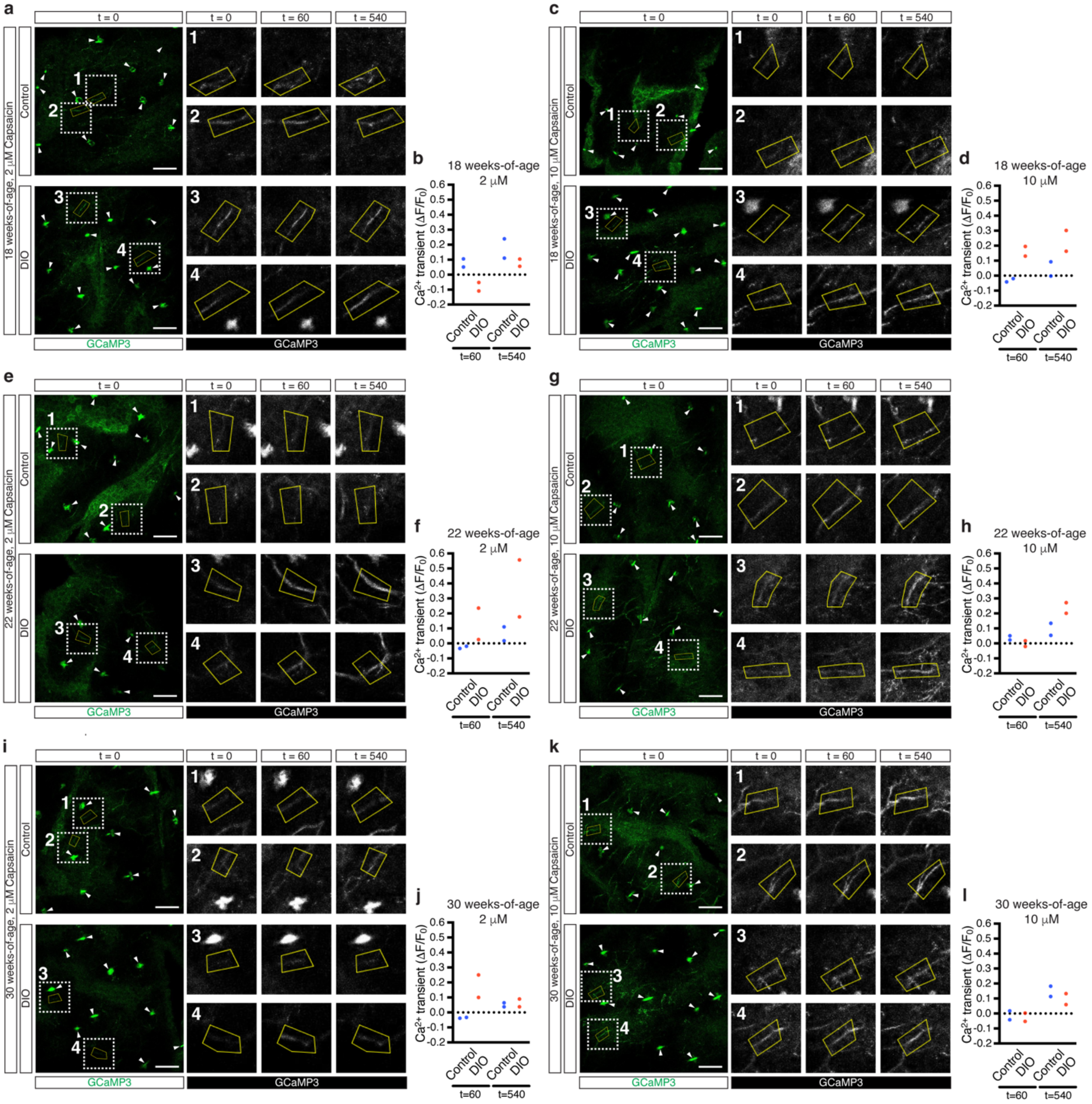
a
b
c
d
e
f
g
h
i
j
k
l
18 weeks-of-age, 2 μM Capsaicin
18 weeks-of-age, 10 μM Capsaicin
22 weeks-of-age, 2 μM Capsaicin
22 weeks-of-age, 10 μM Capsaicin
30 weeks-of-age, 2 μM Capsaicin
30 weeks-of-age, 10 μM Capsaicin
Control
DIO
t = 0
t = 60
t = 540
GCaMP3
18 weeks-of-age
2 μM
10 μM
22 weeks-of-age
30 weeks-of-age
Ca2+ transient (ΔF/F0)
t=60
t=540

**Supplementary Fig. 2. DIO induces sensory hypersensitivity within the epidermis.**

(**a-l**) Representative images showing GCaMP3 expression (green or white) within ear skin explants from *Pirt-GCaMP3* control and DIO mice at 18 weeks-of-age (a-d), 22 weeks-of-age e-h), and 30 weeks-of-age (i-l). Application of 2 μM (a-b, e-f, and i-j) or 10 μM capsaicin (c-d, g-h, and k-l) evoked $Ca^{2+}$ responses in the ear skin explants. Each representative GCaMP3 image at t = 0 is shown on the left (green). The dotted box regions in each image were magnified in the right panels (1-2, control; 3-4, DIO). GCaMP3-expressing axons and their terminals, enclosed by yellow frames, were selected for quantification of fluorescence levels at t = 0, 60, and 540 (white). Representative $Ca^{2+}$ transient ($\Delta F/F_0$) in each selected GCaMP3-expressin axon (1-4) was calculated at t= 60 and 540 (b, d, f, h, j, and l). Note that GCaMP3-expressing hair follicles (arrowheads) were excluded from this analysis. Scale bars: 100 μm.

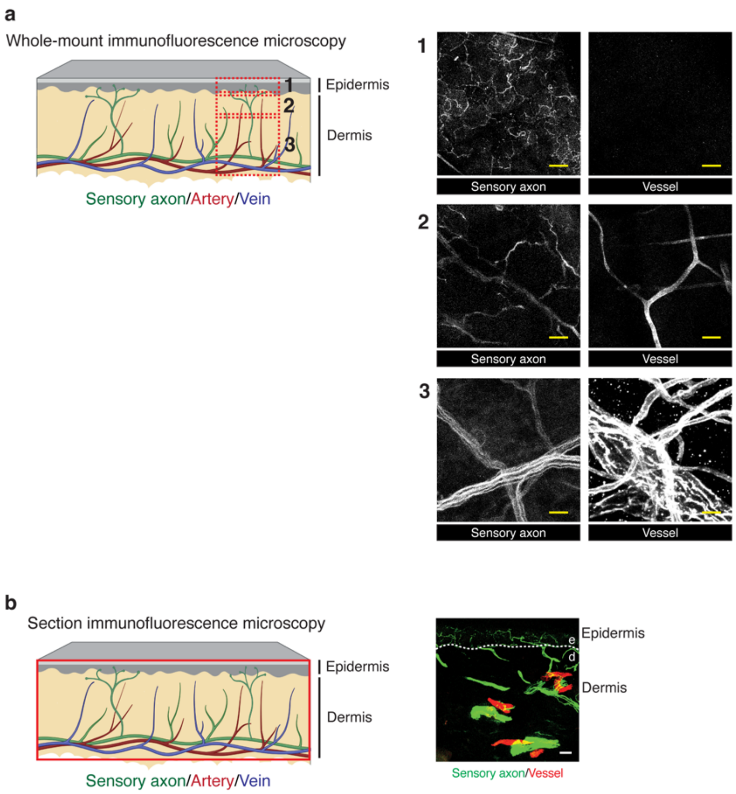


**Supplementary Fig. 3. Schematic diagram of adult ear skin.**

(**a**) A schematic diagram of adult ear skin is shown on the left, with representative whole-mount images displayed on the right. In the epidermis (1), sensory axons and their terminals are present, but capillaries are absent. In the upper dermis (2), both sensory axons and capillaries are present. In the deeper dermis (3), there is co-branching of sensory nerve bundles and large-diameter blood vessels (arteries and veins). (**b**) A schematic diagram of adult ear skin is shown on the left, with a representative section image displayed on the right. As illustrated in (a), the epidermis contains sensory axons and their terminals but lacks capillaries, while the dermis contains both sensory axons and capillaries. Dashed lines indicate the border between the epidermis (e) and the dermis (d). The schematic diagrams were partially created with BioRender.com. Scale bars: 20 μm

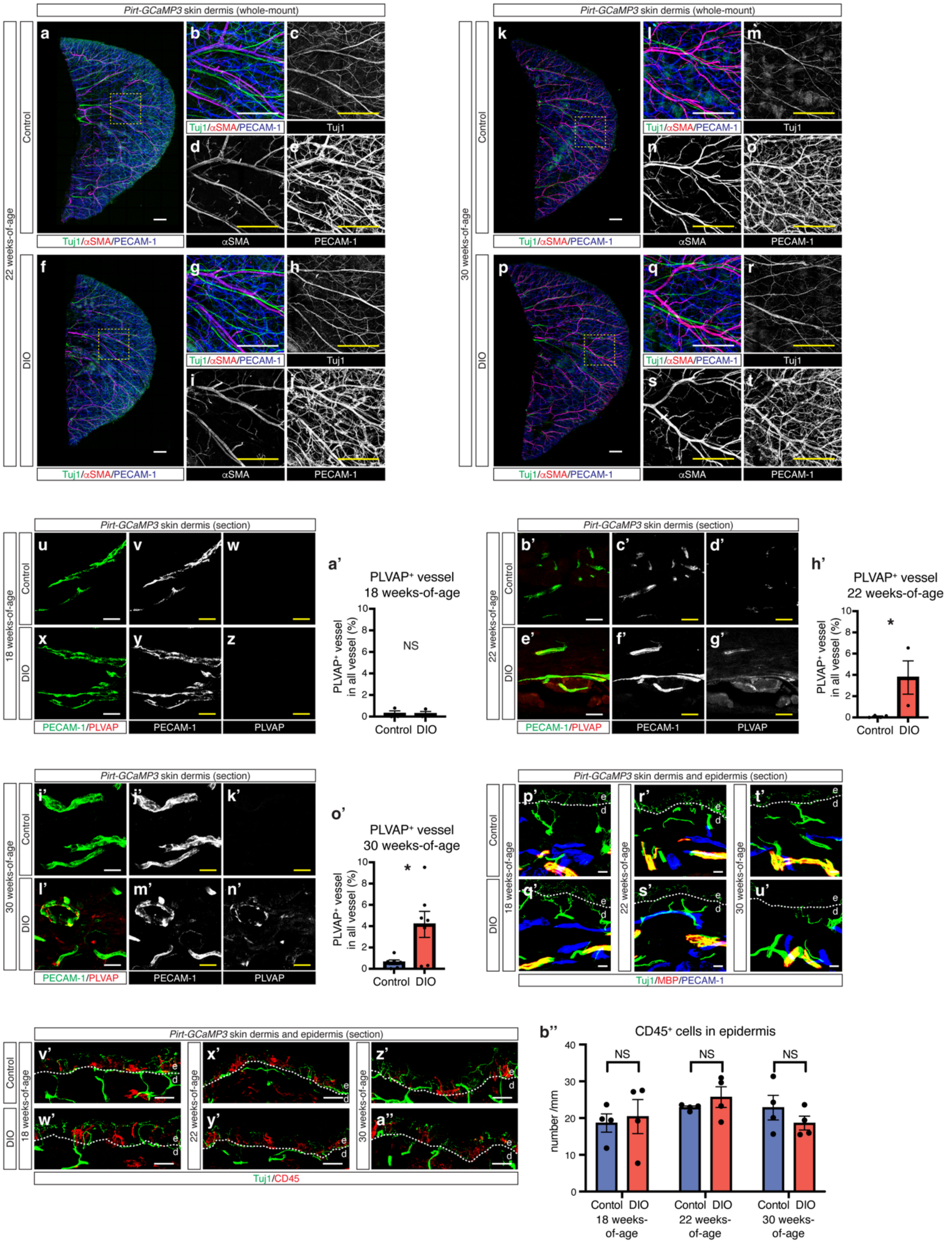
Pirt-GCaMP3 skin dermis (whole-mount)
22 weeks-of-age
30 weeks-of-age
Control
DIO
Tuj1/αSMA/PECAM-1
Tuj1
αSMA
PECAM-1
Pirt-GCaMP3 skin dermis (section)
18 weeks-of-age
PECAM-1/PLVAP
PLVAP
PLVAP⁺ vessel
18 weeks-of-age
PLVAP⁺ vessel in all vessel (%)
NS
Control DIO
PLVAP⁺ vessel
22 weeks-of-age
PLVAP⁺ vessel
30 weeks-of-age
Pirt-GCaMP3 skin dermis and epidermis (section)
Tuj1/MBP/PECAM-1
Tuj1/CD45
CD45⁺ cells in epidermis
number /mm
Contol DIO
18 weeks-of-age
22 weeks-of-age
30 weeks-of-age

**Supplementary Fig. 4. DIO induces vascular permeability in the skin vasculature.**

(**a-t**) Representative whole-mount triple immunofluorescence images of the dermis of ear skin from *Pirt-GCaMP3* control (a-e and k-o) and DIO mice (f-j and p-t) at 22 weeks-of-age (a-j) and 30 weeks-of-age (k-t). Antibodies to the pan-axon/neuron marker neuron-specific class III ß-tubulin (Tuj1, a-b, f-g, k-l, and p-q, green; c, h, m, and r, white), the vascular smooth muscle cell marker αSMA (a-b, f-g, k-l, and p-q, red; d, i, n, and s, white), and the pan-endothelial cell marker PECAM-1 (a-b, f-g, k-l, and p-q, blue; e, j, o, and t, white) were used. The dotted box regions in (a), (f), (k), and (p) were magnified in (b-e), (g-j), (l-o), and (q-t), respectively. Scale bars: 1 mm. (**u-z**, **b'-g'**, and **i'-n'**) Representative double immunofluorescence section images of the dermis of ear skin from *Pirt-GCaMP3* control (u-w, b'-d', and i'-k') or DIO mice (x-z, e'-g', and l'-n') at 18 weeks-of-age (u-z), 22 weeks-of-age (b'-g'), and 30 weeks-of-age (i'-n'). Antibodies to the vascular permeability marker PLVAP (u, x, b', e', i', and l', red; w, z, d', g', k', and n', white) together with PECAM-1 (u, x, b', e', i', and l', green; v, y, c', f', j', and m', white) were used. Scale bars: 20 μm. (**a'**, **h'**, and **o'**) Quantification of $PLVAP^+$ vessels in each image at 18 weeks-of-age (a', N=3 control, N=3 DIO), 22 weeks-of-age (h', N=4 control, N=3 DIO), and 30 weeks-of-age (o', N=6 control, N=7 DIO). The percentage of $PLVAP^+$ vessels in total PECAM-$1^+$ vessels is presented. (**p'-u'**) Representative triple immunofluorescence images of ear skin sections of *Pirt-GCaMP3* control (p', r', and t') and DIO mice (q', s', and u') at 18 weeks-of-age (p'-q'), 22 weeks-of-age (r'-s'), and 30 weeks-of-age (t'-u'). Antibodies to the myelin marker MBP (red) together with Tuj1 (green) and PECAM-1 (blue) were used. Scale bars: 20 μm. (**v'-a''**) Representative double immunofluorescent images of ear skin sections of *Pirt-GCaMP3* control (v', x', and z') and DIO mice (w', y', and a'') at 18 weeks-of-age (v'-w'), 22 weeks-of-age (x'-y') and 30 weeks-of-age (z'-a''). Antibodies to the pan-hematopoietic cell marker CD45 (red) together with Tuj1 (green) were used. Scale bars: 20 μm. (**b''**) Quantification of $CD45^+$ immune cells in the epidermis at 18, 22, and 30 weeks-of-age (N=4 control, N=4 DIO, each stage). The $CD45^+$ immune cell density in the epidermis is calculated and expressed as the number of cells per millimeter of epidermal length. Dashed lines in (p'-a") indicate the border between the epidermis (e) and the dermis (d). Results are shown as the mean ± SEM. *$p<0.05$; NS, not significant ($p > 0.05$). *P* values were determined by the parametric two-tailed *t* test.

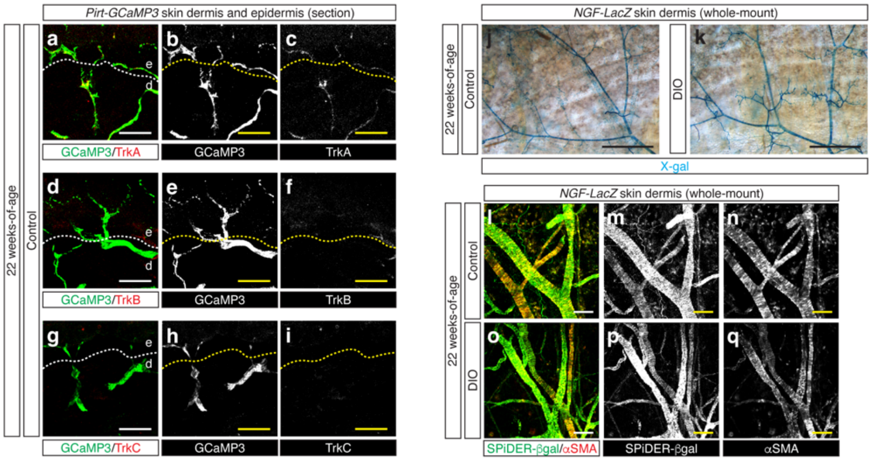


**Supplementary Fig. 5. NGF and its receptor TrkA expression in the ear skin at 22 weeks-of-age.**

(**a-i**) Representative double immunofluorescence images of ear skin sections of *Pirt-GCaMP3* mice at 22 weeks-of-age. Antibodies to GFP for GCaMP3 (a, d, and g, green; b, e, and h, white) together with TrkA (a, red; c, white), TrkB (d, red; f, white), or TrkC (g, red; i, white) were used. Scale bars: 20 µm. (**j-k**) Representative images of whole-mount X-gal staining of the dermis of ear skin from *NGF-LacZ* control (j) and DIO mice (k) at 22 weeks-of-age (blue). Scale bars: 1 mm. (**l-q**) Representative whole-mount double immunofluorescence images of the dermis of ear skin from *NGF-LacZ* control (l-n) and DIO mice (o-q) at 22 weeks-of-age. SPiDER-βgal (l and o, green; m and p, white) together with αSMA antibody (l and o, red; n and q, white) were used. Scale bars: 100 µm.

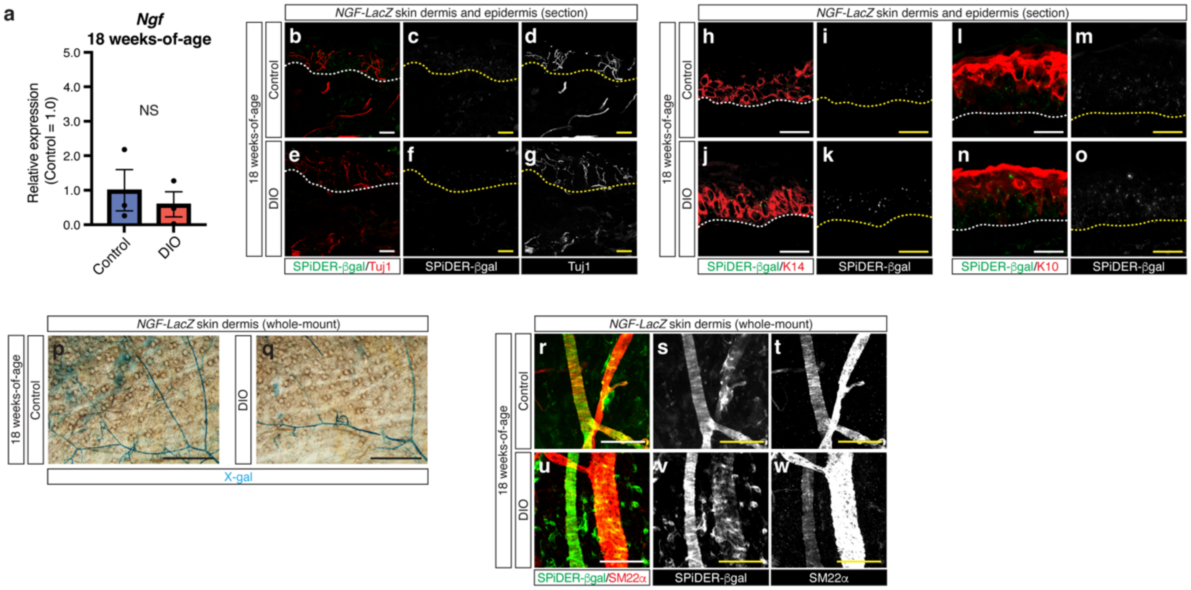


**Supplementary Fig. S6. NGF expression in the ear skin at 18 weeks-of-age.**

(**a**) Relative *Ngf* mRNA expression levels in the epidermis of ear skin from control and DIO mice at 18 weeks-of-age. Expression levels are normalized by those in the epidermis of ear skin from control mice. N=3 in each group. Results are shown as the mean ± SEM. NS, not significant ($p > 0.05$). *P* values were determined by the parametric two-tailed *t* test. (**b-o**) Representative double immunofluorescence images of ear skin sections of *NGF-LacZ* control (b-d, h-i, and l-m) and DIO mice (r-g, j-k, and n-o) at 18 weeks-of-age. SPiDER-βgal (b, e, h, j, l, and n, green; c, f, i, k, m, and o, white) together with antibodies to Tuj1 (b and e, red; d and g, white), K14 (h and j, red), or K10 (l and n, red) were used. Dashed lines indicate the border between the epidermis and the dermis. Scale bars: 20 μm. (**p-q**) Representative images of whole-mount X-gal staining of the dermis of ear skin from *NGF-LacZ* control (p) and DIO mouse (q) at 18 weeks-of-age. Scale bars: 1 mm. (**r-w**) Representative whole-mount double immunofluorescence images of the dermis of ear skin from *NGF-LacZ* control (r-t) and DIO mice (u-w) at 18 weeks-of-age. SPiDER-βgal (r and u, green; s and v, white) together with the vascular smooth muscle cell marker SM22α (r and u red; t and w, white) were used. Scale bars: 100 μm.

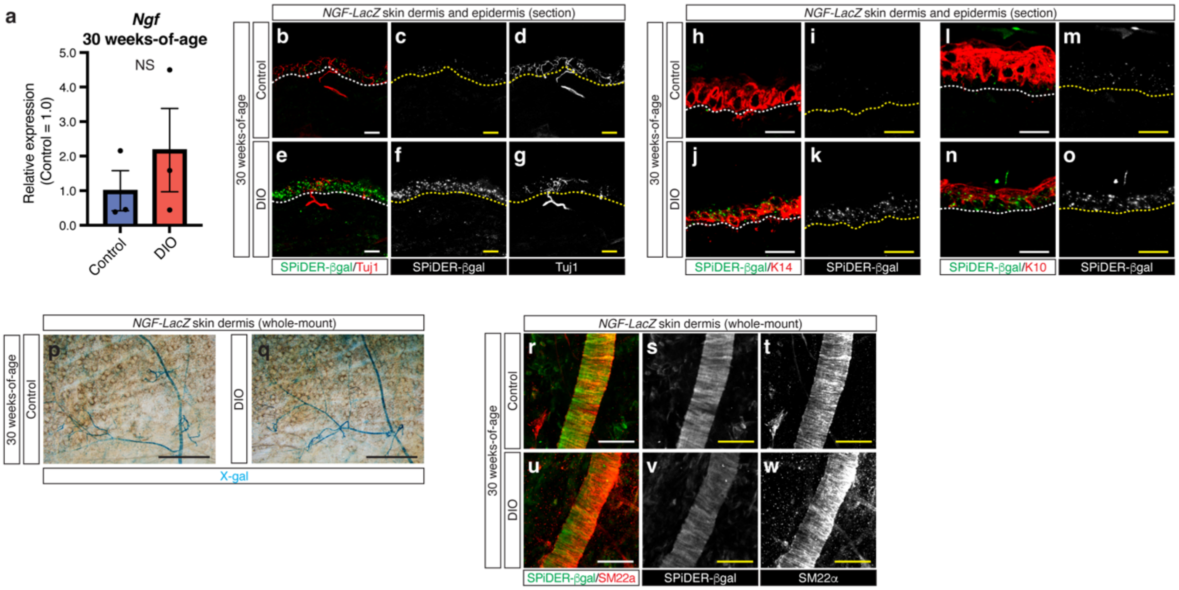


**Supplementary Fig. 7. NGF expression in the ear skin at 30 weeks-of-age.**

(a) Relative *Ngf* mRNA expression levels in the epidermis of ear skin from control and DIO mice at 30 weeks-of-age. Expression levels are normalized by those in epidermis of in the epidermis of ear skin from control mice. N=3 in each group. Results are shown as the mean ± SEM. NS, not significant ($p > 0.05$). *P* values were determined by the parametric two-tailed *t* test. (**b-o**) Representative double immunofluorescence images of ear skin sections of *NGF-LacZ* control (b-d, h-i, and l-m) and DIO mice (e-g, j-k, and n-o) at 30 weeks-of-age. SPiDER-βgal (b, e, h, j, l, and n, green; c, f, i, k, m, and o, white) together with antibodies to Tuj1 (b and e, red; d and g, white), K14 (h and j, red), or K10 (l and n, red) were used. Dashed lines indicate the border between the epidermis and the dermis. Scale bars: 20 µm. (**p-q**) Representative images of whole-mount X-gal staining of the dermis of ear skin from *NGF-LacZ* control (p) and DIO mouse (q) at 30 weeks-of-age. Scale bars: 1 mm. (**r-w**) Representative whole-mount double immunofluorescence images of the dermis of ear skin from *NGF-LacZ* control (r-t) and DIO mice (u-w) at 30 weeks-of-age. SPiDER-βgal (r and u, green; s and v, white) together with SM22α antibody (r and u, red; t and w, white) were used. Scale bars: 100 µm.

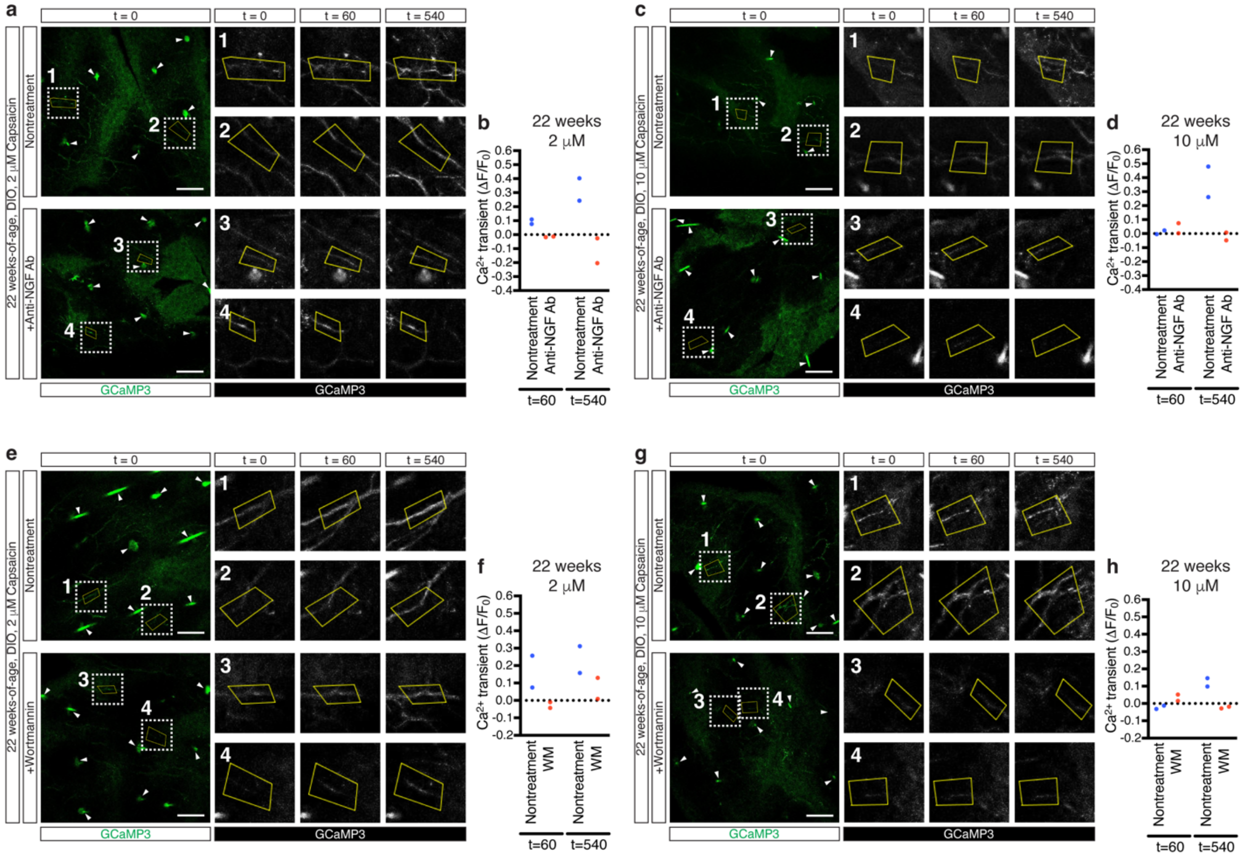


**Supplementary Fig. 8. Inhibition of NGF-TrkA-PI3K signaling suppress hypersensitivity in DIO skin.**

(**a**-**h**) Representative images showing GCaMP3 expression (green or white) within ear skin explants treated with or without anti-NGF neutralized antibody (a-d, nontreatment or +Anti-NGF Ab) and Wortmannin (e-h, Nontreatment or +Wortmannin). Application of 2 μM (a-b, e-f) or 10 μM capsaicin (c-d, g-h) evoked $Ca^{2+}$ responses in the ear skin explants. Each representative GCaMP3 image at t = 0 is shown on the left (green). The dotted box regions in each image were magnified in the right panels (1-2, nontreatment) or (3-4, +Anti-NGF Ab or +Wortmannin). GCaMP3-expressing axons and their terminals, enclosed by yellow frames, were selected for quantification of fluorescence levels at t = 0, 60, and 540 (white). Representative $Ca^{2+}$ transient ($\Delta F/F_0$) in each selected GCaMP3-expressin axon (1-4) was calculated at t= 60 and 540 (b, d, f, h). Note that GCaMP3-expressing hair follicles (arrowheads) were excluded from this analysis.

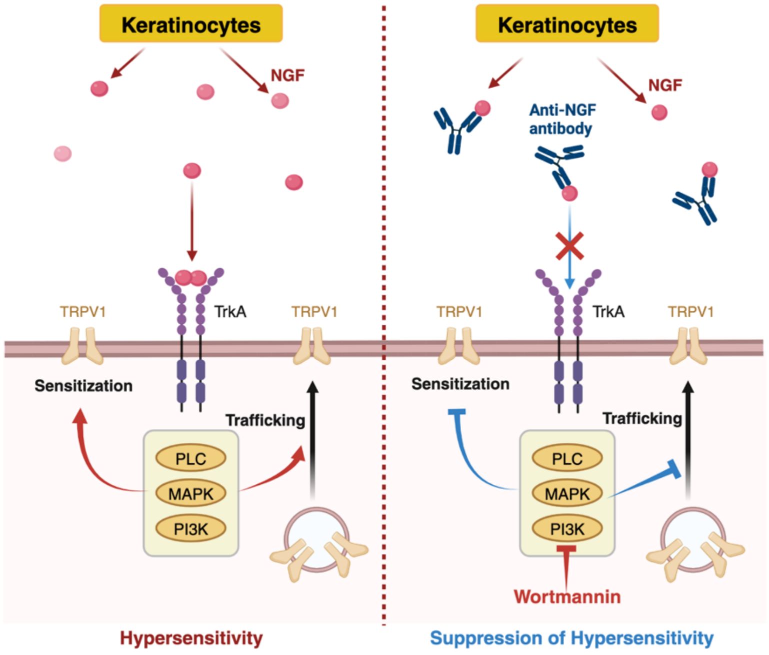


**Supplementary Fig. 9. Keratinocyte-derived NGF sensitizes the peripheral terminals of nociceptive neurons through NGF-TrkA-PI3K signaling within the epidermis of DIO ear skin.**

Keratinocyte-derived NGF sensitizes nociceptive neurons in the DIO skin either through direct sensitization of TRPV1 or by increasing TRPV1 trafficking to the membrane via NGF-TrkA-PI3K signaling. Consequently, treatment of DIO skin with anti-NGF neutralized antibody or Wortmannin treatment leads to the suppression of hypersensitivity.

**Supplementary Table 1. List of quantitative RT-PCR primers for mouse genes.**

| | Forward sequence | Reverse sequence |
|---|---|---|
| *Gapdh* | CTGCACCACCAACTGCTTAG | TCTCATCATACTTGCCAGGT |
| *Ngf* | GTTTTGCCAAGGACGCAGCTTTC | GTTCTGCCTGTACGCCGATCAA |
| *Bdnf* | GGCTGACACTTTTGAGCACGTC | CTCCAAAGGCACTTGACTGCTG |
| *Ntf3* | CTACTACGGCAACAGAGACGCT | GGTGAGGTTCTATTGGCTACCAC |
